# Tumour-derived factors prime microglia for immune checkpoint-mediated T-cell regulation

**DOI:** 10.64898/2026.10.12.758469

**Authors:** Luca D. Schreurs, Jessica Barz, Trong-Hieu Nguyen, Jule Schmedding, Liudmila Lobastova, Patrick Hölker, Tamina Seeger-Nukpezah, Jan-Wilm Lackmann, Jana C. Hollenbeck, Christina Ising, Ulf D. Kahlert, Marco Timmer, Roland Goldbrunner, Volker Neuschmelting, Phuong-Hien Nguyen

## Abstract

Tumour-associated microglia are major regulators of the immune microenvironment in brain malignancies, yet the early mechanisms by which tumour-derived signals shape their immune phenotype remain poorly understood. We used quantitative proteomic analysis on microglia upon tumour exposure and observed extensive remodelling of the microglial proteome following exposure to tumour-conditioned media, indicating that soluble mediators represent the dominant drivers of early microglial activation. Microglia reprogramming was characterized by coordinated upregulation of inflammatory mediators together with proteins involved in antigen presentation and T-cell regulation, including MHC-I, PD-L1 and Galectin-9. Flow cytometric validation using multiple murine and human cell models confirmed the increased expression of PD-L1 and MHC-I on microglia following exposure to tumour-derived conditioned media. Subsequently, tumour-conditioned microglia promoted expression of the inhibitory receptors PD-1 and TIM-3 on T-cells after coculture. Importantly, integration of publicly available single-cell RNA sequencing datasets and cell-cell communication analysis further demonstrated that this immune-regulatory program is recapitulated in microglia from patients with brain tumours. Together, our findings identify an early, soluble factor-driven microglial activation program that precedes direct tumour contact and integrates inflammatory activation with immune checkpoint regulation. These results provide a mechanistic framework for the early establishment of an immunosuppressive brain tumour microenvironment and highlight microglial immune checkpoint pathways as potential targets for therapeutic intervention.

## 1 Introduction

Primary brain tumours and brain metastases (BrM) remain major causes of cancer-related morbidity and mortality (1, 2). Despite considerable advances in neurosurgical techniques, radiotherapy and systemic drug treatments (3), the prognosis of patients with brain malignancy remains poor, with median overall survival rarely exceeding two years (4). Novel immunotherapeutic approaches, which have transformed the treatment of several extracranial malignancies so far, have failed to achieve comparable sustained clinical responses intracranially (2, 3).

Increasing evidence demonstrates that tumour progression and therapeutic resistance are critically shaped by the tumour microenvironment (TME), shifting attention beyond tumour cell-intrinsic mechanisms toward complex cellular interactions within the TME (5). Within the brain TME, tumour-associated macrophages (TAMs) constitute the dominant immune compartment, accounting for approximately 30–40% of non-tumour cells in BrM and an even higher fraction of TME cells in primary brain tumours (5). TAMs comprise both resident microglia and infiltrating monocyte-derived macrophages (MDMs), with their relative contribution dynamically changing during tumour progression. Early tumour lesions are predominantly populated by resident microglia, whereas disruption of the blood-brain barrier during disease progression permits increasing infiltration of MDMs (6, 7, 8).

Resident microglia are increasingly recognized as central regulators of immune responses through extensive interactions with both malignant and immune cell populations. Beyond a sole role as innate immune effector cells (8, 9), microglia actively coordinate inflammatory responses through cytokine secretion, antigen presentation and immune checkpoint signalling.

Effective anti-tumour immunity critically depends on the activation and sustained effector function of tumour-infiltrating T-cells. In brain tumours, however, T-cell activation is constrained by an immunosuppressive microenvironment shaped by resident microglia and infiltrating macrophages, contributing to the limited clinical efficacy of immune checkpoint inhibitors (10).

Although recent transcriptomic and spatial profiling studies have substantially advanced our understanding of tumour-associated microglial phenotypes, comparatively little is known about the functional mechanisms through which tumour-derived factors influence microglial immune responses and their interaction with the lymphoid compartment.

In this study, we therefore investigated how tumour-derived factors modulate microglial phenotypes in the context of brain malignancies, using cellular models of both glioblastoma (GBM) and BrM arising from non-small cell lung cancer (NSCLC). Using proteomics and subsequent validation, we characterized the effects of tumour-derived factors on microglia regarding antigen presentation, immune checkpoint expression and T-cell activation.

## 2 Materials and methods

### 2.1 Isolation of primary murine microglia (pMG)

C57BL6/J wildtype mice were housed under standardized conditions in the CECAD in vivo research facility at the University of Cologne. Pups with mixed genders were sacrificed on postnatal day 1 to 4 (P1-P4) by decapitation and whole brains were removed and transferred into Hank’s Balanced Salt Solution (HBSS). The brains were separated into both hemispheres and cerebellum, the olfactory bulbs and the meninges were carefully removed under a microscope. Following three washes in HBSS, brain tissue was dissociated enzymatically with 0.25% trypsin for 10 min at 37°C. Enzymatic digestion was stopped by adding microglia medium consisting of DMEM-GlutaMAX supplemented with 10% heat-inactivated FBS and 1% penicillin/streptomycin. DNase was used for resuspension to obtain a homogenous single cell solution. Cells were seeded into flasks precoated with 0.1% poly L-lysin and incubated at 37°C in a humidified incubator containing 5% CO2 and 92% relative humidity. After two to three days, cells were washed with warm Dulbecco’s phosphate-buffered saline (DPBS) and microglia medium supplemented with 10% L929 conditioned media was added. Cultures were maintained for an additional five to seven days after which primary mouse microglia (pMG) were harvested by tapping. pMG were transferred to RPMI medium supplemented with 10% FBS and 1% penicillin/streptomycin for subsequent experiments. Microglia were harvested up to three times.

### 2.2 Treatment of microglia and macrophages with conditioned media (CM)

Tumour cells were seeded at a density of 5 x 10^5^ cells/ml in RPMI-1640 and allowed to attach overnight. After 24 hours, culture medium was replaced with fresh medium, and CM was harvested following an additional 24-hour incubation period. CM was centrifuged at 900g for 10 minutes at room temperature (RT) and subsequently filtered using 0.2 µm filters.

Microglia and macrophages were seeded in 6-well plates at a density of 1 x 10^6^ per well in RPMI-1640 and were allowed to attach for 24 hours before replacing the medium with freshly harvested CM. After another 24 hours, CM was removed and cells were washed twice with DPBS. Cells were detached using Accutase and processed for flow cytometric analysis. All antibodies used are listed in Table 1. Medium-treated controls were included to determine baseline expression levels.

### 2.3 Proteomics analysis

Primary mouse microglia were seeded at a density of 1 × 10^6^ cells per well in 6-well plates containing microglia medium. A total of 12 wells were prepared, corresponding to three experimental conditions with four biological replicates, each covering untreated control (UT), direct co-culture (dCC), and conditioned medium (CM).

KPC were used to generate both tumour spheroids and CM. For spheroid formation, 1 × 10^4^ KPC in 100 µL complete RPMI (cRPMI) medium were seeded into each well of an ultra-low attachment 96-well plate and centrifuged at 300 g for 5 s. In parallel, 1 × 10^4^ KPC in 1 mL RPMI were seeded into each well of a 24-well plate for conditioned medium production. The following day, spheroids were centrifuged at 300 g for 5 s.

For CM, KPC cultures were washed twice with DPBS and transferred from cRPMI to serum-free RPMI medium for 24 hours. Simultaneously, pMG cultures were switched from MG medium to serum-free RPMI. One KPC spheroid was transferred to each microglia well assigned to the direct co-culture condition. Prior to transfer to microglia cultures, undiluted CM was collected after 24 h of serum-free conditioning and clarified by sequential centrifugation at 300 g for 5 min and 1,000 g for 10 min, followed by filtration through a 0.2 µm Millipore filter.

For dCC, KPC spheroids were removed from co-cultures after 24 hours by microscope-assisted microdissection. Microglia were washed with DPBS, detached by scraping, collected by centrifugation, washed once in PBS, and transferred to 1.5 mL microcentrifuge tubes.

Microglia cell pellets were resuspended in 10 µL SP3 lysis buffer (5% SDS in 1× PBS) and lysed by repeated pipetting. Chromatin was fragmented using a Bioruptor sonicator for 10 minutes with alternating 30 s on/off cycles at room temperature. Dithiothreitol (DTT) was added to a final concentration of 5 mM, and samples were incubated at 55°C for 30 minutes. Chloroacetamide (CAA) was subsequently added to a final concentration of 40 mM, and samples were incubated for 30 min at room temperature in the dark. Afterwards, samples were digested using LysC and trypsin (both Promega) following a modified SP3 protocol by foregoing peptide cleanup on day two and instead directly load samples onto SDB-RPS Stage Tips for purification and analysed after elution.

Samples were analyzed by the CECAD Proteomics Facility on an Orbitrap Eclipse mass spectrometer (Thermo Scientific) coupled to an Evosep One (Evosep) running in 30 SPD (samples per day) setup with the recommended chromatographic column. MS1 scans were acquired from 385 m/z to 1015 m/z at 30k resolution. Maximum injection time was set to 118 ms and the AGC target to 250%. MS2 scans ranged from 400 m/z to 1000 m/z and were acquired at 30k resolution with a maximum injection time of 54 ms, an AGC target of 1000%, and were acquired in 20 x 30 m/z staggered windows shifted by 15 m/z for each second cycle, resulting in nominal 12 m/z windows after deconvolution using ProteoWizard (Chambers 2012). All scans were stored as centroid.

Samples were afterwards analyzed using Spectronaut 19. Files were converted into htrm followed by a direct-DIA search against a Swissprot mouse canonical database (UP589, downloaded 04/01/25) with settings matching acquisition parameters. Afterwards, analysis of results was performed in Perseus 1.6.15 (Tyanova 2016) by filtering for data completeness in replicate groups, FDR-controlled T-tests, and 1D enrichment analysis.

### 2.4 Integration of single-cell RNA sequencing datasets and cell-cell interaction analysis

Raw data were downloaded from GEO, the raw data already contained gene names, barcodes, and count matrix. Single cell data analysis was performed using the Seurat v5.0 suite (11). We removed cells whose % mitochondrial genes > 10%, ambient RNA contamination was estimated by decontX from the celda package (12). Dimensional reduction (PCA, UMAP) and clustering were done by Seurat functions with default hyper-parameters. We integrated 4 NSCLC samples and integrate 2 GBM samples separately. Both datasets were integrated by CCA methods implemented in Seurat, default hyper parameters. Finally, we inferred cell-cell interaction using CellChat (13).

## 3 Results

### 3.1 Exposure to tumour-derived factors suppresses homeostatic microglial programs while promoting inflammatory activation, metabolic, and functional remodelling of pMG

To investigate early changes in microglial phenotype following exposure to tumour cells, pMG were cultured either with CM derived from murine KPC or in direct coculture (dCC) with a KPC spheroid prior to label-free quantitative proteomic analysis (Figure 1A, Supplementary Figure 1A). Both conditions induced remodelling of the pMG proteome compared with untreated controls. Tumour CM treatment resulted in 358 significantly differentially expressed proteins (DEPs), whereas dCC only led to 15 (Figure 1B-C, Supplementary Figure 1B). In contrast, no DEPs were identified between the direct coculture and conditioned medium conditions (Figure 1D), indicating that tumour-derived soluble factors account for the majority of the early proteomic changes observed following microglial exposure to KP cells. In total, both conditions induced 559 DEPs in pMG (Figure 1E, Supplementary Figure 1C).

**Figure 1:**
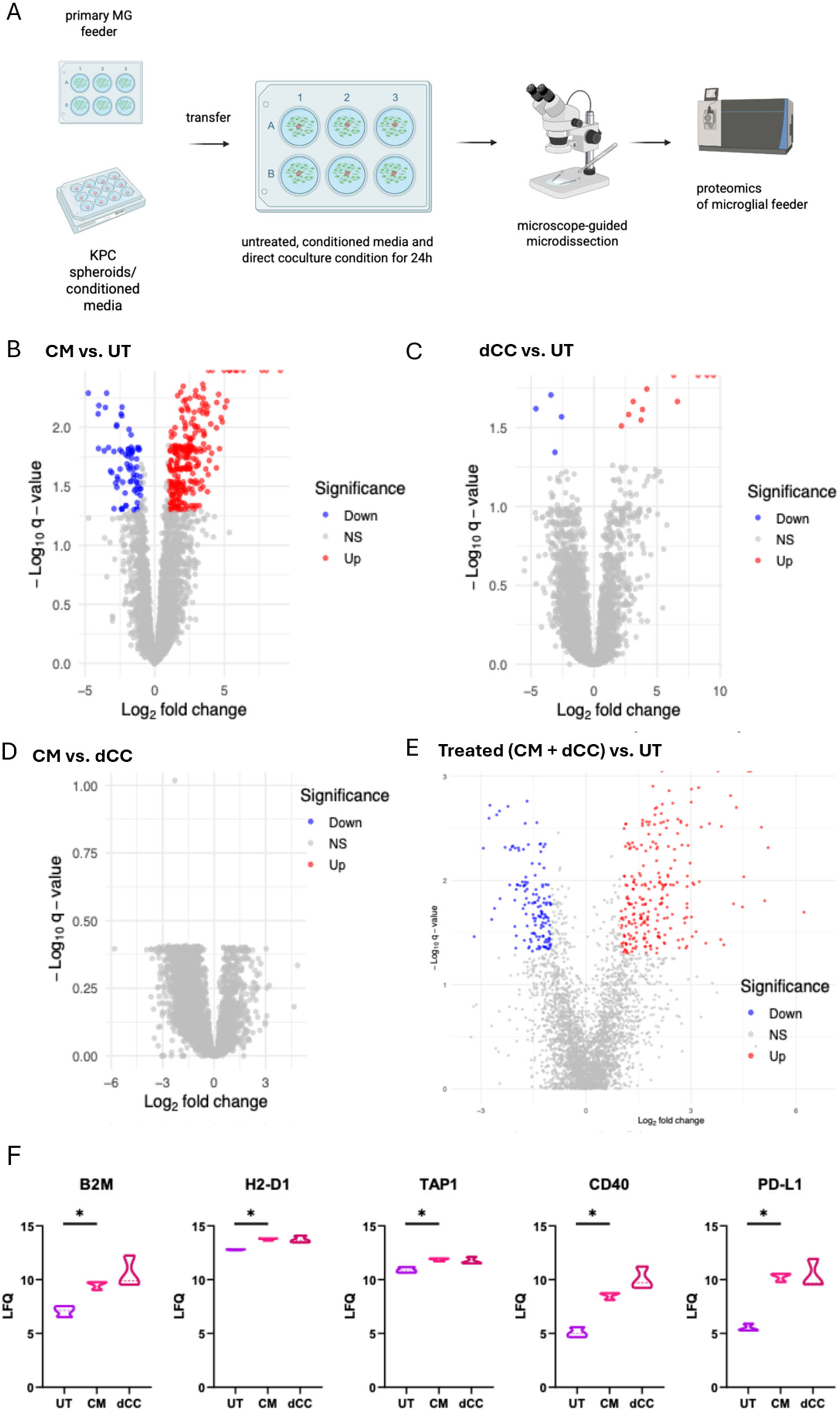
Proteomic analysis of primary microglia (pMG) exposed to KPC conditioned media (CM) and KPC spheroids. (A) Experimental setup: pMG (feeder layer) were either treated with KPC CM, were cocultured with KPC spheroids or were left untreated (UT) for 24 hours. Microglial feeder cells were isolated by microscope-guided microdissection and subjected to proteomic analysis. (B-E) Volcano plots comparing proteomic profiles between conditions: CM vs. UT (B) , dCC vs. UT (C), Treated (CM vs. dCC) vs. UT (D) and CM vs. dCC (E). Significantly upregulated proteins are shown in red, significantly downregulated proteins are shown in blue and non-significant proteins in grey. (F) Violin plots showing label free quantification (LFQ) intensities of PD-L1, H2-D1, TAP1 and CD40 and B2M in microglia across UT, CM and dCC conditions. *p < 0.05.

Exposure to tumour CM resulted in pronounced induction of proteins associated with inflammatory and innate immune activation. The most strongly increased proteins included the scavenger receptor MARCO, the immunometabolic enzyme ACOD1, prostaglandin E synthase (PTGES), the chemokine CCL5, lipocalin-2 (LCN2), and the activation marker CD69 (Supplementary Figure 2A). Additionally, inflammatory mediators, including IL1A, IL1B, TNF, GPR84 and CLEC4E, were likewise increased following tumour exposure (Supplementary Figure 2B). Notably, other proteins classically associated with disease-associated microglia showed reduced abundance upon exposure to tumour-derived factors. These included complement proteins (C1QA, C1QC) and osteopontin (SPP1) (Supplementary Figure 2C). Reduced expression was particularly pronounced for the TAM-receptor ligand GAS6 (Supplementary Figure 2C). Further, other proteins linked to immune-regulation showed decreased abundance upon tumour exposure, including the folate receptor FOLR2 and the glycoprotein receptor CD200R1 (Supplementary Figure 2C). Together, these changes indicate that soluble factors released by KPC induce a broad inflammatory activation program that shows overlap but also distinct differences to classical disease-associated microglia.

**Figure 2:**
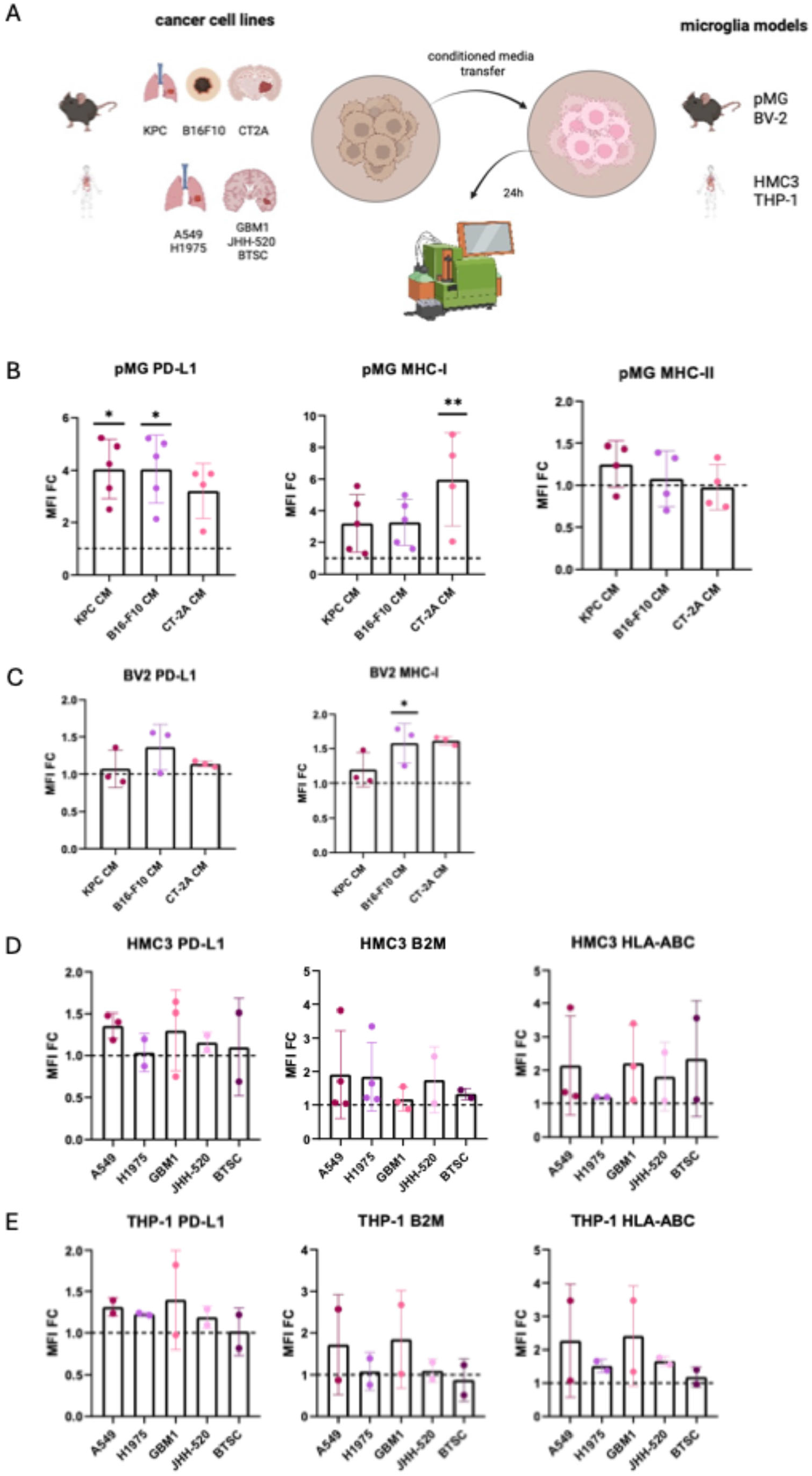
Tumour-derived CM upregulated immune checkpoint and antigen presentation-associated proteins across microglia models. Experimental setup: Murine microglia (pMG, BV2) and human microglia (HMC3, THP-1) were respectively treated with CM from murine cancer cell lines (KPC, B16-F10, CT-2A) and human cancer cell lines (A549, H1975, GBM1, JHH-520, BTSC) for 24 hours and analyzed via flow cytometry. Mean fluorescence intensity (MFI) fold-change (FC) of PD-L1, MHC-I and MHC-II in pMG following exposure to KPC, B16-F10 and CT-2A CM. MFI FC of PD-L1 and MHC-I in BV2 cells following exposure to KPC, B16-F10 and CT-2A CM. *p < 0.05. MFI FC of PD-L1, B2M and HLA-ABC in HMC3 cells following exposure to A549, H1975, GBM1, JHH-520, BTSC CM. MFI FC of PD-L1, B2M and HLA-ABC in THP-1 macrophages following exposure to A549, H1975, GBM1, JHH-520, BTSC CM. The dotted line indicates baseline MFI measured in the untreated control cells. Each dot represents an independent experiment with three replicates. *p < 0.05, **p < 0.01.

In line, the phenotypic changes were accompanied by an increased abundance of several classical signalling proteins implicated in myeloid cell activation and inflammatory responses, including SRC, NOTCH2, CEBPB, JAK2, and HCK (Supplementary Figure 3A). Proteins involved in immunometabolism and cellular redox homeostasis including SLC7A11, SLC20A1, SLC2A1, HMOX1, SOD2, SRXN1, GSS, ARG2, and SQSTM1 displayed increased abundance following tumour CM exposure (Supplementary Figure 3B). Exposure to tumour CM also altered the abundance of proteins involved in phagocytosis, cell adhesion, and extracellular matrix remodelling. Increased expression of MSR1, PLAUR, ICAM1, PILRA, PILRB, FSCN1, MMP9, MMP13, and PLXNA1 were observed (Supplementary Figure 3C), demonstrating remodelling of pathways involved in environmental sensing, cellular motility, and interactions with the surrounding microenvironment.

**Figure 3:**
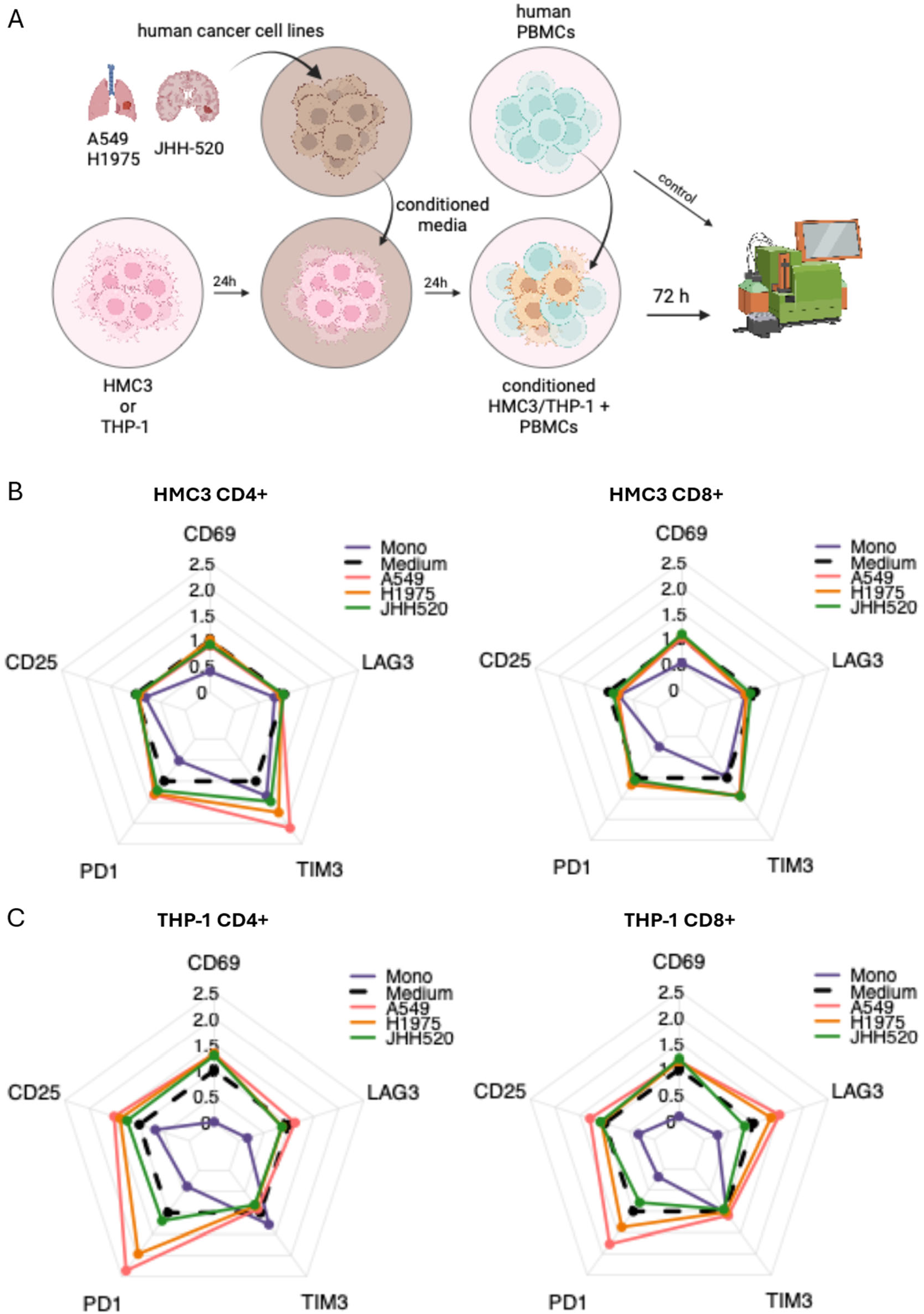
Tumour-conditioned HMC3 and THP-1 cells modulated T-cell activation and inhibitory receptor expression. Experimental setup. CM from human cancer cell lines (A549, H1975, JHH-520) was applied to HMC3 cells or THP-1 macrophages for 24 hours. The conditioned HMC3 and THP-1 cells were then cocultured with human PBMCs, followed by flow cytometric analysis after 72 hours. Radar plots showing expression of activation and immune checkpoint markers (CD69, CD25, LAG3, TIM3, PD1) on CD4+ and CD8+ T-cells after coculture with HMC3 cells, which were pretreated with A549, H1975 and JHH-520 CM. Comparison to monoculture (Mono) and medium controls.Radar plots showing expression of activation and immune checkpoint markers (CD69, CD25, LAG3, TIM3, PD1) on CD4+ and CD8+ T-cells after coculture with THP-1 cells, which were pretreated with A549, H1975 and JHH-520 CM. Comparison to monoculture (Mono) and medium controls.

### 3.2 Tumour-conditioning promotes the acquisition of an antigen-presenting and T-cell-interacting phenotype of microglia

Interestingly, proteomic analysis revealed coordinated regulation of proteins involved in adaptive immune communication. Components of the major histocompatibility complex class I antigen presentation machinery, including B2M, H2-D1, and TAP1, were increased following tumour CM exposure (Figure 1F). In parallel, the co-stimulatory molecule CD40 and the immune checkpoint ligand PD-L1 (CD274) were markedly upregulated (Figure 1F). These data demonstrate that exposure to tumour CM induced a coordinated proteomic program in primary mouse microglia characterized by enhanced antigen presentation, co-stimulatory and checkpoint signalling, and increased expression of proteins involved in communication with adaptive immune cells.

To validate whether microglia indeed acquired an immune reactive phenotype upon tumour exposure beyond the KPC-pMG model, different microglia models were exposed to CM generated from multiple tumour models representing primary and metastatic brain malignancies, in both mouse and human systems (Figure 2A).

Consistent with the proteomic analysis, all murine tumour CM significantly increased PD-L1 expression in pMG compared with untreated controls (Figure 2B). Similarly, MHC-I expression was markedly elevated, with CT-2A-conditioned media inducing the strongest response (Figure 2B). In contrast, MHC-II expression remained largely unchanged across conditions (Figure 2B). The immortalized murine microglia cell line BV2 was also exposed to these murine tumour CM, which similarly induced a consistent increase in PD-L1 and MHC-I expression (Figure 2C), albeit in smaller magnitude than pMG. B16-F10-derived conditioned media produced the strongest PD-L1 induction, whereas CT-2A-conditioned media elicited the greatest increase in MHC-I expression.

Using the human microglia cell line HMC3 exposed to CM from NSCLC and glioma models (Figure 2A), we observed consistently upregulated PD-L1 compared with untreated controls (Figure 2D). While the magnitude of PD-L1 induction was moderate, the response was reproducible across all tumour conditions. HLA-ABC expression was likewise increased, with glioblastoma-derived conditioned media producing the strongest induction, indicating that distinct tumour entities differentially modulate antigen-presentation pathways in human microglia. Finally, we examined THP-1-derived macrophages, a commonly used surrogate model for human microglia. Similar to HMC3 cells, tumour CM increased PD-L1, B2M and HLA-ABC expression relative to untreated controls (Figure 2E). However, THP-1 macrophages exhibited greater variability between tumour conditions and an overall less pronounced response than HMC3 cells, suggesting cell type-specific differences in sensitivity to tumour-derived immunomodulatory signals.

Collectively, these findings validate the proteomic profiling of pMG and demonstrate that tumour-derived soluble factors consistently induce immune checkpoint and MHC-I-associated antigen-presentation pathways across murine and human models.

### 3.3 Tumour-conditioned microglia differentially modulate the expression of immune checkpoint molecules on T-cells

To investigate whether tumour-conditioned changes in microglia can subsequently influence T-cell phenotypes, HMC3 and THP-1 cells were pretreated with CM derived from lung adenocarcinoma A549 and H1975, as well as the GBM cell line JHH-520 before coculture with healthy donor PBMCs. Expression of the activation markers CD69 and CD25 together with the inhibitory receptors PD-1, TIM3 and LAG-3 were analysed in CD4+ and CD8+ T-cell subsets after three days of coculture (Figure 3A).

Upon coculture with tumour-primed HMC3 microglia, CD4+ T-cells displayed a modest increase in PD-1 expression and a strong increase in TIM-3 expression, particularly after coculture with HMC3 cells primed with A549- and H1975-derived CM (Figure 3B). A similar but less pronounced increase in PD-1 and TIM-3 expression could also be observed in the CD8+ T-cell subset. The activation markers CD69 and CD25, as well LAG-3 expression showed only minor changes (Figure 3B).

After coculture with tumour-conditioned THP-1 macrophages, both CD4+ and CD8+ T-cells exhibited increased CD69 and CD25 expression (Figure 3C). PD-1 expression was likewise increased, particularly following coculture with lung tumour-conditioned THP-1 macrophages. In contrast to the HMC3 cell line, LAG-3 expression increased while TIM-3 expression remained similar to monoculture controls (Figure 3C).

Overall, pretreatment of both HMC3 and THP-1 cells with tumour-derived CM consistently altered the expression of T-cell activation markers and inhibitory receptors in subsequent coculture, with some distinct temporal cell line-dependent differences in inhibitory receptor expression such as TIM-3 and LAG-3.

### 3.4 Patient datasets identify conserved TAM-to-T-cell immune checkpoint signalling

To determine the relevance of the identified microglial T-cell regulatory interactions, whether they can also be observed in patients, we analysed available published single-cell RNA sequencing (scRNAseq) datasets from NSCLC BrM (four integrated datasets, GSE131907) and GBM (two integrated datasets, GSE182109). Consistent with the distinct immune landscapes of these malignancies (Schreurs et al., 2025), NSCLC BrM exhibited substantially greater lymphoid cell infiltration and immune cell diversity than GBM, whose microenvironment was predominantly composed of TAMs (Figure 4A). Cell–cell communication analysis using CellChat (13) predicted extensive intercellular signalling networks in both tumour types (Figure 4B). Given the central role of T-cell subsets in anti-tumour immunity and immunotherapy, we focused our CellChat analysis on T-cells as signal receiver cells. As expected, MHC-I-mediated antigen presentation to cytotoxic T-cells was predicted to originate from multiple cell populations, including tumour cells and different TAM subsets (Figure 4C).

**Figure 4:**
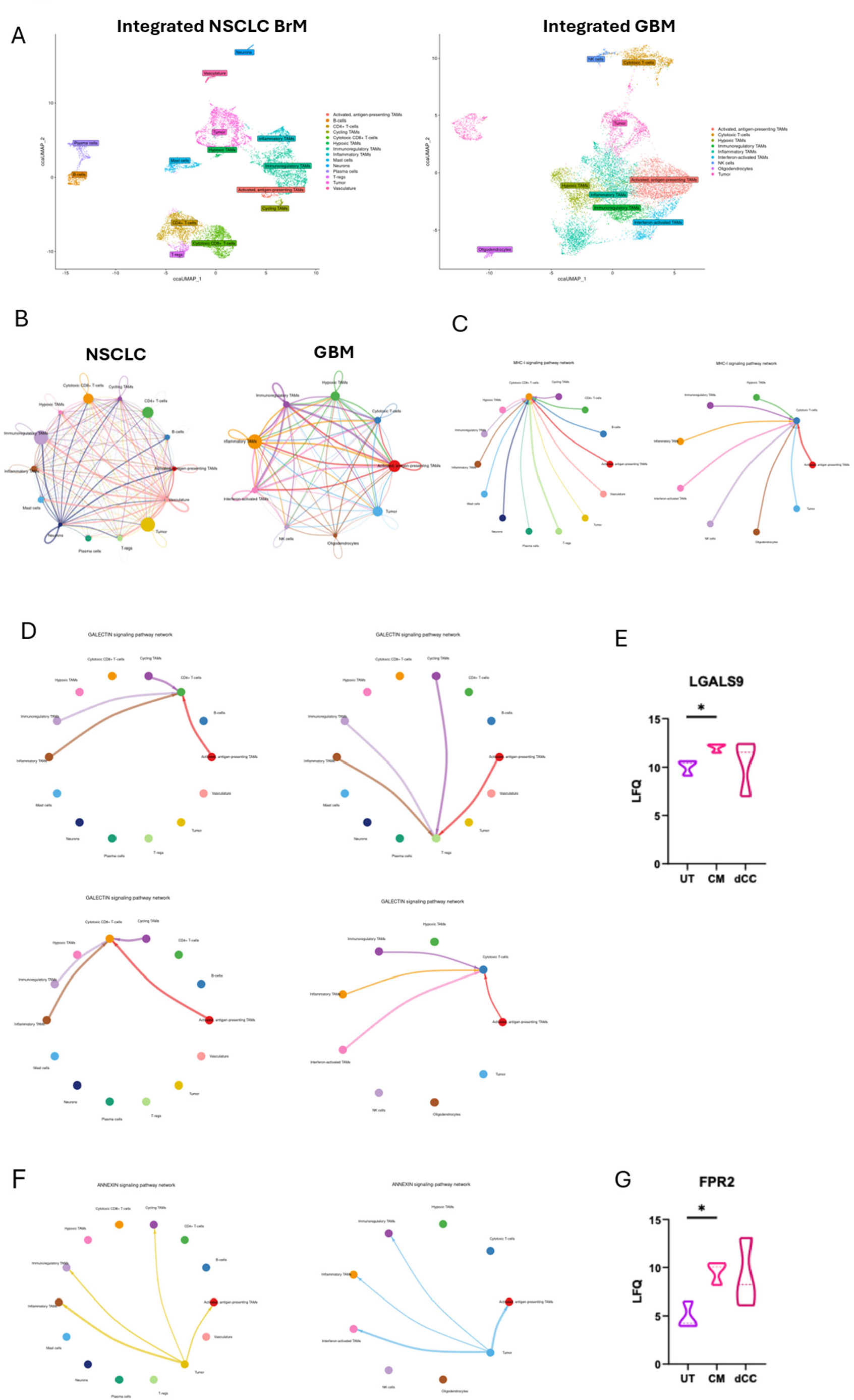
Integration of patient single-cell RNA sequencing datasets and CellChat analysis identified conserved cell-cell communication. (left) UMAP of integrated single-cell data from NSCLC BrM showing annotated cell populations including tumour cells, TAM subsets (activated, antigen-presenting; cycling; hypoxic; immunoregulatory and inflammatory), T-cell subsets (CD4+ T-cells; cytotoxic CD8+ T-cells; T-regs), B-cells, plasma cells, mast cells, neurons and vasculature. (right) UMAP of integrated single-cell data from GBM showing annotated cell populations including tumour cells, TAM subsets (activated, antigen-presenting; hypoxic; immunoregulatory; inflammatory and Interferon-activated), cytotoxic T-cells, NK cells and oligodendrocytes. Circle plot showing the aggregated weighted interaction network across all identified cell populations in NSCLC BrM and GBM.Circle plot showing the MHC-I received signal in cytotoxic CD8+ T-cells in (left) NSCLC BrM and (right) the MHC-I received signal in cytotoxic T-cells in GBM.Circle plot showing the GALECTIN received signal in CD4+ T-cells, in T-regs, in cytotoxic CD8+ T-cells all in NSCLC BrM and in cytotoxic T-cells in GBM.Violin plot showing LFQ intensity of LGALS9 (Galectin-9) in primary microglia across UT, CM and dCC conditions.
Circle plot showing the Annexin signal from (left) NSCLC BrM and (right) GBM sent to TAMs.Violin plot showing LFQ intensity of FPR2 in primary microglia across UT, CM and dCC conditions. *p < 0.05.

We next examined whether the immune checkpoint-mediated TAMs-to-T-cell interactions could also be detected in the scRNAseq datasets. CellChat analysis identified LGALS9 (Galectin-9) signalling as the predominant immune checkpoint pathway, which originated exclusively from TAMs in both NSCLC BrM and GBM, consistent with our in vitro observations identifying microglia as a major source of T-cell inhibitory signals (Figure 4D). Within both tumour entities, Galectin-9 signalling primarily originated from immunoregulatory, inflammatory and activated antigen-presenting TAM subsets. Additional signalling was observed from cycling TAMs in NSCLC BrM and interferon-activated TAMs in GBM, whereas hypoxic TAMs did not contribute in either. Galectin-9 signalling was received by cytotoxic T-cell populations in both tumour types and additionally by CD4+ T-cells and regulatory T cells in NSCLC BrM. In line with this, Galectin-9 showed increased abundance in our proteomic profiling of pMG after exposure to tumour CM (Figure 4E). The identification of this TAM-derived Galectin-9 signalling in two distinct brain tumour datasets provides independent support for our experimental observations and extends the relevance of the Galectin-9–TIM-3 axis to the human disease setting.

Finally, to gain insight into possible mechanisms by which tumours educate TAMs toward an immunosuppressive phenotype, we next analysed tumour-to-TAM communication. Comparison of tumour-to-TAM communication revealed limited conservation between NSCLC BrM and GBM. The only pathway consistently identified in both tumour entities was Annexin signalling, mediated by tumour-derived ANXA1 and Humanin (MT-RNR2) interacting with FPR2 expressed on TAMs (Figure 4F). Supporting this prediction, our proteomic profiling of pMG demonstrated increased FPR2 abundance following exposure to tumour-derived CM (Figure 4G).

## 4 Discussion

Our study sought to identify conserved mechanisms of tumour-induced immune remodelling and to define how these changes influence adaptive immune responses within the brain tumour microenvironment. By integrating primary murine microglia, immortalized murine and human microglial models, we demonstrated that tumour-derived soluble factors are sufficient to induce a rapid and reproducible immunoregulatory reprogramming of microglia, which is characterized by coordinated upregulation of proteins involved in antigen presentation and T-cell regulation, including MHC-I, PD-L1 and Galectin-9. Functionally, tumour-conditioned microglia promoted the expression of inhibitory receptors such as PD-1 and TIM-3 on T-cells, suggesting an exhaustion phenotype. Importantly, these responses were not only observed across diverse murine and human cell lines, but also reflected in patient-derived scRNAseq datasets, suggesting a conserved response of microglia to brain tumour-derived signals.

### 4.1 Tumour-educated microglia represent an early transitional activation state

Our findings further refine current concepts of microglial activation in BrM. Previous studies have demonstrated that BrM-associated TAMs only partially overlap with classical disease-associated microglia (DAM), instead of acquiring inflammatory and immune-regulatory phenotypes characterized by enhanced cytokine signalling and antigen presentation (14, 15, 16). Our proteomic data is consistent with these observations, revealing robust induction of inflammatory mediators together with proteins involved in antigen presentation and immune regulation. However, our data also suggests that early tumour-induced microglial activation may not fully conform to the discrete TAM states proposed from patient datasets. Gonzalez et al. (17) described two principal TAM populations in brain metastases, an immunomodulatory population expressing proteins including SPP1 and HLA molecules, and a second inflammatory population enriched for cytokines and chemokines with lower HLA expression. In contrast, microglia exposed to tumour-derived factors in our study simultaneously acquired inflammatory features together with increased expression of antigen presentation machinery and inhibitory immune regulators. Rather than representing previously described states, these data may suggest an intermediate activation phenotype that combines features of both programs. Several factors may explain these differences. First, many published single-cell studies analysed all TAMs collectively and did not clearly distinguish resident microglia from monocyte-derived macrophages, despite their distinct developmental origins and activation programs. Second, our experimental systems model the earliest stages of microglial activation following exposure to tumour-derived soluble factors, whereas patient specimens represent chronically established tumour microenvironments with prolonged reciprocal interactions between tumour cells and infiltrating immune populations have already occurred. Finally, species-specific differences between murine models and human patient samples may additionally contribute to the observed variation. Together, these considerations suggest that the phenotype identified in our study may represent an early transitional state preceding the more specialized TAM populations observed in established tumours.

### 4.2 Tumour-derived factors establish and immunoregulatory microglia phenotype

The pronounced induction of antigen presentation machinery observed in our study is consistent with previous reports describing increased MHC-I expression in GBM-associated microglia (18). Likewise, induction of PD-L1 following exposure to glioma-CM has previously been reported in murine MG (19), while PD-L1 expression has been documented in human BrM from multiple primary tumour entities and correlates with intratumoral inflammation (20, 21). Our data extend these observations by demonstrating that coordinated induction of antigen presentation proteins and immune checkpoint ligands occurs rapidly following exposure to tumour-derived soluble factors, indicating that these phenotypic changes arise early during tumour-microglia interactions.

### 4.3 Tumour-educated microglia drive immune checkpoint-mediated T-cell suppression

The functional consequences of this phenotype are supported by both our study and previous observations. TAMs spatially associate with exhausted T-cells in both GBM and BrM (16, 22), while PD-L1 expression on TAMs correlates with PD-1 expression on infiltrating T-cells in BrM (17). Consistent with these findings, conditioned microglia in our study promoted expression of PD-1 and TIM-3 or LAG-3 on T-cells. Beyond cancer, microglial PD-L1 has been shown to directly suppress CD8+ T-cell responses during viral CNS infection and experimental autoimmune neuroinflammation (23, 24, 25), supporting a conserved immunoregulatory function of this pathway within the central nervous system.

In addition, our data identified Galectin-9 as a potential contributor to microglia-mediated T-cell regulation. TIM-3 expression on tumour-infiltrating T-cells has repeatedly been reported in brain tumours (19, 26), and recent single-cell analyses identified recurrent Galectin-9–TIM-3 interactions within the BrM microenvironment, particularly between microglia and PD-1-positive CD8+ T-cells (27). Moreover, PD-1 and TIM-3 signalling appear to cooperate functionally, with Galectin-9 supporting the persistence of exhausted PD-1+TIM-3+ T-cells (28). Altogether, these findings suggest that multiple inhibitory checkpoint pathways may be activated simultaneously during early microglial reprogramming. This concept is of clinical interest given the development of TIM-3-directed immunotherapies, which have shown encouraging early clinical activity in models of brain malignancies (29, 30).

Supporting this immunosuppressive role, depletion of microglia after tumour establishment reduced metastatic burden while decreasing T-cell exhaustion and increasing cytotoxic T-cell infiltration in BrM models (14). Mechanistically, CXCL10-mediated recruitment of PD-L1-positive microglia contributed to T-cell dysfunction. Conversely, Evans et al. (15) reported increased metastatic growth following microglial depletion before tumour engraftment, accompanied by impaired T-cell priming. Rather than representing contradictory functions, these findings may reflect distinct temporal roles of microglia during metastatic progression. Notably, the depletion strategies differed between studies, whereas Guldner et al. (14) targeted microglia after tumours had become established and microglial education had already occurred, Evans et al. (15) depleted microglia before metastatic seeding. Our results support a model in which microglia initially contribute to immune surveillance but are subsequently reprogrammed by tumour-derived signals into actively promoting immune evasion.

### 4.4 Conclusion and outlook

Several limitations should still be considered. First, although our data establish that tumours can induce immunoregulatory microglial phenotypes, the upstream mediators responsible for this transition remain to be determined. Multiple cytokines implicated in brain tumours, including IFN-γ, IL-6, TGF-β and IL-33, have previously been reported to induce PD-L1 expression in microglia and therefore represent plausible candidates (31, 32). We found that Annexin signalling represented the only conserved tumour-to-TAM pathway across both tumour entities. Given the concomitant upregulation of FPR2 in tumour-conditioned pMG observed in our proteomic analysis, these data raise the possibility that ANXA1/Humanin–FPR2 signalling represents a shared early mechanism by which distinct brain tumours educate resident myeloid cells toward an immunoregulatory phenotype, as previously suggested for breast cancer BrM (33). Future studies should determine the relative contribution of these pathways to microglia functions and investigate whether they converge on common transcriptional regulators. Second, our experimental model only captures a predominantly unidirectional interaction from tumour cells to microglia, whereas continuous reciprocal communications with various T-cell populations in vivo exert further modifications to microglial phenotypes. For example, interactions with activated T-cells have been proposed to promote more reactive microglial states in human brain tumours (26). Incorporating these multicellular interactions will therefore be important for defining the dynamic evolution of microglial phenotypes during tumour progression.

In summary, our study identifies tumour-derived soluble factors as sufficient to initiate an early immunoregulatory reprogramming of microglia, characterized by coordinated induction of antigen presentation machinery and immune checkpoint ligands, which in turn regulates T-cell activities. By promoting PD-1 and TIM-3 expression on T-cells, tumour-reprogrammed microglia may contribute to the progressive establishment of an immunosuppressive TME, highlighting microglial immune checkpoint pathways as potential therapeutic targets for restoring anti-tumour immunity.

## Statements

### Data availability statement

Data are available upon request to the corresponding author.

### Ethics statement

PBMCs were collected at the University Hospital of Cologne after written and informed consent according to the Declaration of Helsinki and with Institutional Review Board approvals at the University of Cologne no. 19-1559, no. 21-1317, and no. 19-1438_1.

Mouse experiments were approved by the state authorities of North Rhine-Westphalia, Germany (Landesamt für Natur-, Umwelt-und Verbraucherschutz Nordrhein-Westfalen (LANUV)), approval no. UniKoeln_Anzeige §4.22.008.

## Conflict of interest

All authors declare no conflict of interest.

### Generative AI statement

The authors declare that generative AI was not used in the creation of this manuscript.

