## Supplementary Information for "Tumour-derived factors prime microglia for immune checkpoint-mediated T-cell regulation"

**Supplementary Material**

**Supplementary Methods**

**1. Cell culture**

All cell lines were cultured in RPMI-1640 or DMEM supplemented with 10% fetal bovine serum (FBS) and 1% penicillin/streptomycin. Cells were maintained at 37°C in a humidified incubator containing 5% CO_2_ and 92% relative humidity. Culture medium was routinely refreshed according to standard cell culture protocols and cells were regularly monitored for morphology and confluency.

HMC3 and THP-1 cell lines served as human microglia and macrophage models. Tumour-derived conditioned media (CM) were harvested from human lung adenocarcinoma cell lines A549 and H1975, as well as the human GBM cell lines GBM1, JHH-520 and BTSC (1).

Murine experiments included primary microglia (pMG) isolated from C57BL/6 mice, the microglial cell line BV2, the immortalized *Kras*-mutant *Trp53*-deficient lung adenocarcinoma cells KPC (2), the melanoma cell line B16-F10 and the glioblastoma cell line CT-2A.

**2. Differentiation of THP-1 monocytes into macrophages**

THP-1 cells were seeded in 6-well plates at a density of 1 x 10^6^ cells per well in RPMI-1640. Differentiation into macrophage-like cells was induced by addition of phorbol 12-myristate 13-acetate (PMA) at a final concentration of 100 ng/ml. After 24 hours of stimulation, cells were washed thoroughly with phosphate-buffered saline (PBS) to remove residual PMA and cultured in fresh RPMI-1640. Successful differentiation was assessed by adherence and morphological changes using light microscopy (3).

**3. Isolation of peripheral blood mononuclear cells (PBMCs)**

Whole blood was diluted with sterile PBS supplemented with 2% FBS (1:1 dilution). For density gradient separation, 15 ml of Pancoll was added to SepMate^TM^ Isolation Tubes (STEMCELL^TM^ Technologies, Cat #85450. The diluted blood was carefully layered onto the density gradient medium without disturbing the interface and centrifuged at 1200 g for 10 minutes at RT with the brake turned off. Following centrifugation, the plasma layer was carefully removed using a pipette and the PBMC-containing interphase was collected and transferred into a new tube. PBMCs were washed twice using PBS with 2% FBS and were centrifuged at 350 g for 10 minutes at 4 °C between washing steps. Residual erythrocytes were removed by incubation with 4 ml of ACK Lysing Buffer (Gibco). PBMCs were washed again to remove the remaining lysis buffer and were resuspended in freezing medium containing 50% complete RPMI (cRPMI), 40% FBS, 10% dimethyl sulfoxide (DMSO). Cells were stored at -150 °C until further use.

**4. Cocultures of HMC3/THP-1 with PBMCs**

HMC3 cells or THP-1-derived macrophages were seeded into 6-well plates at a density of 5 x 10^3^ cells per well and were incubated for 24 hours to allow stable attachment prior to experimental treatment. Cells were subsequently incubated with tumour-derived CM for 24 hours. Following CM exposure, medium was replaced completely with fresh culture medium and PBMCs were added at a density of 2 x 10^5^ cells per well, corresponding to a coculture ratio of 1:40. Cocultures were maintained to enable direct cell-cell interactions and T-cell activation. PBMC monocultures served as baseline controls. T-cell activation was assessed after 72 hours by flow cytometric analysis of activation and immune checkpoint markers. At each timepoint, 100 µl of cell suspension was collected and processed for flow cytometry analysis. CD3+ and CD4+ or CD8+ T-cells were analyzed.

**5. Flow cytometry**

Flow cytometry was performed using the MACSQuant X (Miltenyi Biotec) flow cytometer as previously described (3). All reagents and antibodies are listed in the Supplementary Table.

**6. Statistical analysis**

All experimental conditions were tested in technical triplicates for each time point. Each experiment was repeated as an independent biological replicate to ensure reproducibility. All data was normalized to the respective medium control prior to statistical analysis by appropriate non-parametric statistical tests to compare experimental conditions using GraphPad Prism and R-Studio. A P value < 0.05 was considered statistically significant, while correcting for multiple testing when needed.

Figures 1A, 2A, 3A were created with Biorender.

**Supplementary Table:** List of antibodies and reagents used for flow cytometric analysis

| **Anti-** | **Fluorochromes** | **Supplier** | **Cat. & Lot.** |
| --- | --- | --- | --- |
| Annexin V | VioBlue | BioLegend | Cat. 640917  Lot. B320623 |
| CD25 (IL-2R) | Brilliant Violet 510 | BioLegend | Cat. 302640  Lot. B366218 |
| PD-1 (CD279) | Vio Bright 515 | Miltenyi Biotec | Cat. 130-120-392  Lot. 5211200786 |
| CD3 | PE | Miltenyi Biotec | Cat. 130-113-139  Lot. 5241009726 |
| CD8 | PerCP Vio700 | Miltenyi Biotec | Cat. 130-110-682  Lot. 52511077845 |
| CD69 | PE-Cy7 | BioLegend | Cat. 310911  Lot. B292790 |
| LAG-3 | APC | BioLegend | Cat. 369212  Lot. B311711 |
| TIM-3 (CD366) | APC-Cy7 | BioLegend | Cat. 345025  Lot. B363500 |

**Supplementary Figures**

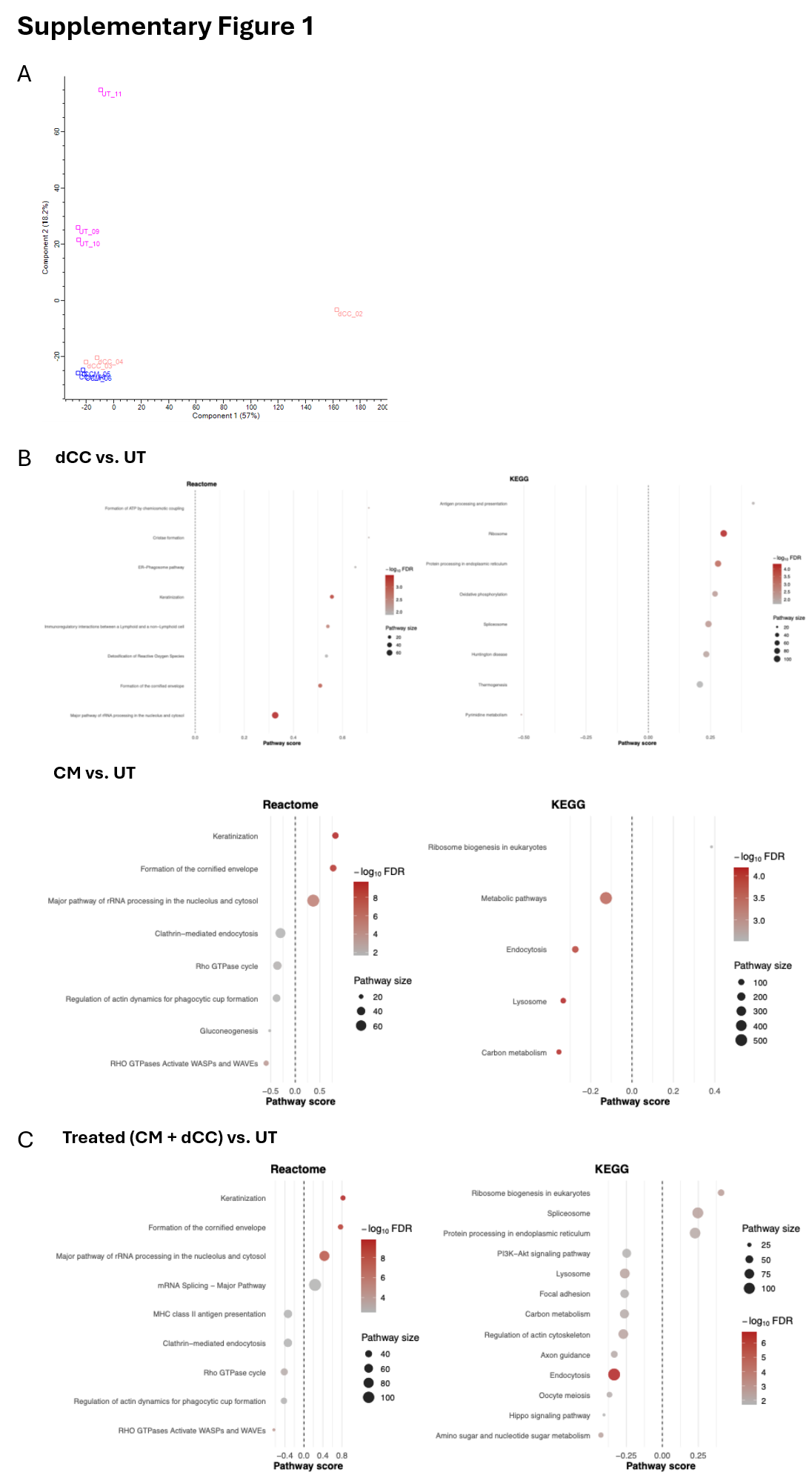

**Supplementary Figure 1**

(A) PCA of proteomic analysis of pMG showing UT, CM and dCC conditions

(B) Reactome and KEGG pathway analysis dCC vs. UT and CM vs. UT. Dot size represents the pathway size.

(C) Reactome and KEGG pathway analysis for Treated (CM+dCC) vs. UT. Dot size represents the pathway size.

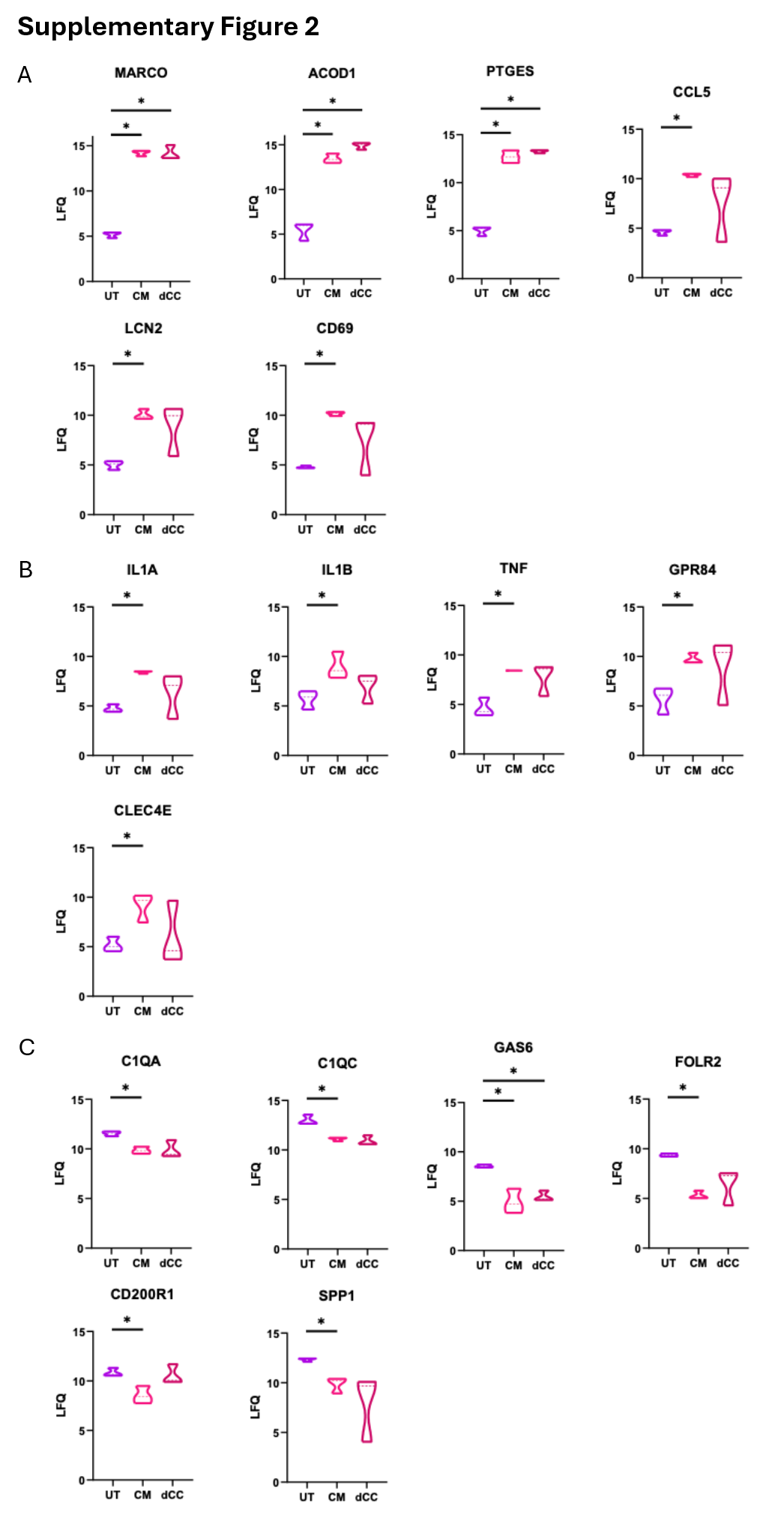

**Supplementary Figure 2**

(A) Violin plots showing LFQ intensities of activation-associated proteins MARCO, ACOD1, PTGES, CCL5, LCN2 and CD69 in pMG across UT, CM and dCC conditions.

(B) Violin plots showing LFQ intensities of proinflammatory mediators IL1A, IL1B, TNF, GPR84 and CLEC4E in pMG across UT, CM and dCC conditions.

(C) Violin plots showing LFQ intensities of microglia markers C1QA, C1QC, GAS6, FOLR2, CD200R1 and SPP1 in pMG across UT, CM and dCC conditions.

*p < 0.05 **p < 0.01

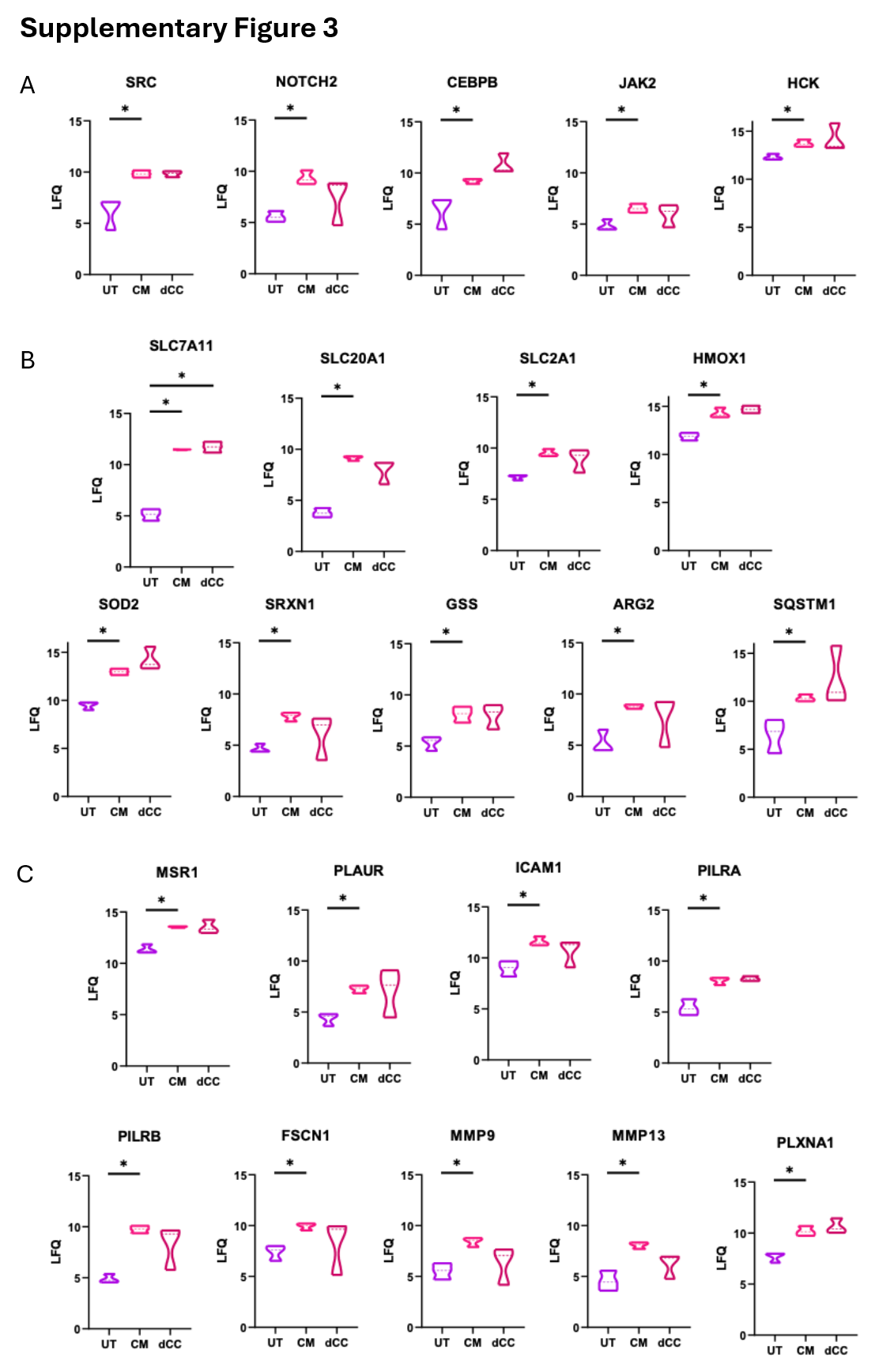

**Supplementary Figure 3**

(A) Violin plots showing LFQ intensities of signaling proteins SRC, NOTCH2, CEBPB, JAK2, HCK in pMG across UT, CM and dCC conditions.

(B) Violin plots showing LFQ intensities of metabolic and stress response proteins SLC7A11, SLC20A1, SLC2A1, HMOX1, SOD2, SRXN1, GSS, ARG2 and SQSTM1 in pMG across UT, CM and dCC conditions.

(C) Violin plots showing LFQ intensities of antigen-presentation related proteins MSR1, PLAUR, ICAM1, PILRA, PILRB, FSCN1, MMP9, MMP13 and PLXNA1 in pMG across UT, CM and dCC conditions.

*p < 0.05 **p < 0.01
